# *PRESSED FLOWER* works downstream of *ASYMMETRIC LEAVES 2* to affect sepal flatness in *Arabidopsis*

**DOI:** 10.1101/2024.09.30.615753

**Authors:** Ruoyu Liu, Zeming Wang, Xi He, Heng Zhou, Yiru Xu, Lilan Hong

## Abstract

The development of flattened organs such as leaves and sepals is essential for proper plant function. While much research has focused on leaf flatness, little is known about how sepals achieve flat organ morphology. Previous study has shown that in Arabidopsis an *ASYMMETRIC LEAVES 2* (*AS2*) gene mutation *as2-7D* causes ectopic *AS2* expression on the abaxial sepal epidermis, which leads to growth discoordination between the two sides of sepals, resulting in outgrowth formation on abaxial sepal epidermis and sepal flatness disruption. Here we report that the *PRESSED FLOWER* (*PRS*) works downstream of *AS2* in affecting sepal flatness. Genetic analysis showed that *PRS* mutations suppressed the outgrowth formation on the abaxial sepal epidermis in *as2-7D* mutant. Through tracking the *PRS* expression dynamics at a cellular resolution throughout the early developmental stages in WT and *as2-7D* sepals, we found that on the abaxial epidermis of *as2-7D* sepals, ectopic *AS2* expression up-regulated *PRS* expression, leading to the epidermal outgrowth initiation. AS2 affected PRS activity on multiple levels: AS2 activated *PRS* expression through direct binding to *PRS* promoter region; AS2 also physically interacted with PRS. Our study highlights the complex interplay between AS2 and PRS in modulating sepal flatness.

## INTRODUTION

During development, plant organs display a myriad of complex shapes in three dimensions: cylindrical shoots, spherical or oblong fruits, lobed anthers, leaves and petals that can be flat, twisting, bending, waving, cup-shaped, tubular, etc (Tena, 2024). Flat morphology is a key structural form in plant organs, with flat leaves being the most common example (Tsukaya, 2005; Sandalio *et al*., 2016). The development of three asymmetrical axes, an adaxial-abaxial axis, a medio–lateral axis, and a proximal–distal axis, plays an crucial role in the transformation of the leaf blade from a radial primordium to a flattened structure during leaf growth and development (Ichihashi and Tsukaya, 2015; Nakayama *et al*., 2022). Extensive molecular genetic studies have identified a regulatory network for the establishment of adaxial-abaxial polarity in leaves, involving auxin and abaxial- and adaxial-promoting genes (Lin *et al*., 2007; Shi *et al*., 2017). How the proximal-distal polarity is established remains largely unknown (Du *et al*., 2018), while the establishment of the medio-lateral polarity (from the midrib to the margin) depends on the adaxial-abaxial polarity (Waites and Hudson, 1995).

The leaf blade exhibits anisotropic growth during morphogenesis, with cell division occurring primarily perpendicular to the medio-lateral axis, which results in flattened leaves (Du *et al*., 2018). Growth along the medio-lateral axis is contingent on the activity of leaf meristems (Nardmann and Werr, 2013; Ichihashi and Tsukaya, 2015; Guan *et al*., 2017). Although leaves exhibit determinate growth without typical anatomical features of meristematic tissue, there is transient leaf meristematic activity allowing the expansion of leaf blade (Alvarez *et al*., 2016). In *Arabidopsis thaliana* (*Arabidopsis*), the transient leaf meristematic activity, reflected in part by the expression of *WUSCHEL-RELATED HOMEOBOX3* (*WOX3*) / *PRESSED FLOWER* (*PRS*), is progressively confined to the entire marginal region in young leaves and further confined to the proximal marginal region in older leaves (Alvarez *et al*., 2016). *PRS* and *WUSCHEL-RELATED HOMEOBOX1* (*WOX1*) act redundantly in the marginal region (also called the middle domain) between the adaxial and abaxial domains and are critical for leaf blade outgrowth (Vandenbussche *et al*., 2009; Nakata *et al*., 2012). *PRS* and *WOX1* work through coordinately regulating the proliferation of *WOX*-expressing cells and surrounding cells (Nakata *et al*., 2012). When ectopic expression of *WOX1* and *PRS* occurs in the abaxial domain of leaf primordia, outgrowths form on the abaxial surface of leaves (Nakata *et al*., 2012).

Sepals, the exterior floral organs of most flowering plants, also exhibit a flattened morphology, which helps to protect the development of internal flower organs (Roeder, 2021). While many studies have focused on the flattened morphology of leaves, relatively few studies have explored on the morphogenesis of flattened organs based on sepals. In fact, *Arabidopsis* sepal serves as an excellent model for studying organ morphogenesis due to its simplicity, accessibility, and reproducibility of morphogenesis. Sepals are the most peripheral organ of the flower, making them easy to manipulate and observe (Zhu *et al*., 2020). The final size of the *Arabidopsis* sepal is only about 1 mm^2^, and the entire morphogenesis of living sepals can be imaged using high magnification microscopes (Hong *et al*., 2016). Plant growth, development, and morphogenesis are continuous processes that occur in both space and time. Dynamic observation of these biological processes has been a significant technical challenge in plant developmental biology. The use of time-lapse live imaging combined with data processing and analysis tools in *Arabidopsis* sepals allows for simultaneous temporal and spatial observation of sepal size, shape, and curvature, as well as tracking of sepal cell growth and division dynamics (Hong *et al*., 2017; Robinson *et al*., 2018). In addition, based on transcriptomic analysis of floral organs, only 13 genes were found to be specifically expressed in sepals; in other words, most genes expressed in sepals are also expressed in other organs (Wellmer *et al*., 2004). Therefore, insights gained from studying *Arabidopsis* sepals can be broadly applied to understanding the morphogenesis of other plant lateral organs.

In our previous work, we identified a mutant with abnormal sepal morphology, *as2-7D*, which has an unbalanced growth on the abaxial-adaxial surfaces of the sepals, resulting in numerous folds and outgrowths on the abaxial surface. The folds comprise ridges and invaginations, whereas the outgrowths, which are pointed epidermal projections, are typically located at the end of the folds. The *as2-7D* mutant sepal epidermis produces outgrowths due to conflicting growth directions and unequal epidermal stiffness. The mutant phenotype was found to be caused by a mutation in the *AS2* gene by map-based cloning (Yadav *et al*., 2024). The *AS2* gene encodes a transcription factor that contains a *LATERAL ORGAN BOUNDARIES (LOB)* domain (Iwakawa *et al*., 2002; Lin *et al*., 2003).The *AS2* promoter region has a KANADI (KAN) transcription factor binding site, and the binding of KAN to the *AS2* promoter represses *AS2* expression (Wu *et al*., 2008). A point mutation occurred at the KAN binding site in *as2-7D* genome, which altered the expression pattern of the *AS2* gene. The wild type (WT) *AS2* promoter is active on the adaxial surface of the organ, while the mutant *AS2* promoter drives expression on both the abaxial and adaxial surfaces of the sepals (Yadav *et al*., 2024). Although the effect of ectopic *AS2* expression on sepal flatness have been explored in detail on the cellular level, the molecular mechanisms underlying *AS2*’s function in regulating sepal flatness remain not fully understood.

In this study, to further explore the genes involved in controlling sepal flatness, we did a suppressor screen of *as2-7D’*s sepal phenotype and obtained a suppressor mutant with reduced sepal epidermal outgrowths. The *PRS* gene was identified as the mutant gene of the suppressor. Using live imaging we provided a detailed depiction of the *PRS* expression pattern at a cellular resolution throughout the early developmental stages of WT and *as2-7D* sepals. We found that on the abaxial epidermis of *as2-7D* sepals, the ectopic expression of *AS2* tends to retain *PRS* expression and keeps it to linger on the rising outgrowth, which initiates epidermal outgrowths initiation. Furthermore, AS2 interacts with PRS at the protein level and promotes *PRS* expression at the transcriptional level, contributing to the formation of outgrowths in *as2-7D* sepals. Our work highlights the complex interplay between AS2 and PRS in modulating sepal flatness.

## METHOD

Plant materials and growth conditions Landsberg *erecta* (L*er*) ecotype plants is the WT in this study. The *as2-7D* mutant was previously identified by our research group in a screen (Yadav *et al*., 2024). *as2-7D* has a G to A point mutation at 1484 bp upstream of the *AS2* gene promoter, resulting in ectopic expression of the *AS2* gene. ^60^Co-γ rays were used to mutagenize *as2-7D* seeds (radiation dose rate of 10 Gy·min^-1^, radiation dose of 600 Gy). In the M2 generation of the self-pollinated progeny from the M1 plants, a mutant showing significant suppression of the sepal outgrowth phenotype was identified and designated as *as2-7D suppressors* of *as2-7D sepal phenotype 1* (*as2-7D ssp1*). Crossing *as2-7D ssp1* with the WT resulted in the isolation of the single mutant *ssp1* (*prs-3*) in the F2 generation. The *prs-4* mutant (T-DNA insertion line *SALK_127850*) was obtained from the Arabidopsis Biological Resource Center (ABRC). The *as2-5D* mutant has been previously described (Wu *et al*., 2008) and was ordered from ABRC (CS67863). Plants were grown at 22°C under 16-hour light/8-hour dark conditions, with a light intensity of 12,000 Lux.

### QTL-seq

In this study, *as2-7D* × *as2-7D ssp1*was hybridized, and individual plants with phenotypes of *as2-7D* and *as2-7D ssp1* were selected from the F2 population to construct two mixed offspring pools with extreme phenotypes for QTL-seq analysis (Takagi *et al*., 2013). The Δ SNP index was obtained by subtracting the SNP index of *as2-7D ssp1* phenotype pool with the SNP index of *as2-7D* phenotype pool. The Δ SNP index is plotted with the genome position as the horizontal axis and the Δ SNP index as the vertical axis to obtain the Δ SNP index map. In the plot, the region where the peak or valley is located in the map is the candidate region for the actual mutation site. QTL-seq analysis showed that there were differences in SNP index on chromosome 2 between the two separate populations, and the *SSP1* suppression gene was preliminarily located in the genomic region of 9 Mb-16 Mb on chromosome 2.

### Flower staging

Flower staging was based on Smyth et al (Smyth *et al*., 1990). The main observations in this study were on flowers at around stage 6 when outgrowth formation initiated in *as2-7D* sepals.

### Sepal architecture

Flowers are enclosed by four sepals, of which the out-most sepal is located away from the meristematic tissue and is called the abaxial/outer sepal, while the sepal that is diametrically opposed to the outer sepal and oriented toward the meristematic tissue is called the adaxial/inner sepal. The remaining two sepals located on either side are the lateral sepals. For a detailed description, see Roeder (Roeder, 2021).

### Scanning electron microscopy

Sepals at stage 14 were fixed overnight in 2.5% glutaraldehyde solution-0.2M sodium phosphate buffer (pH 7.0) at 4 □. The fixed samples were washed three times with 0.1 M sodium phosphate buffer (pH 7.0) for 15 minutes each time. The samples were washed with 0.1 M phosphate buffer solution (pH 7.0) and dehydrated through a graded ethanol series (30%, 50%, 70%, 80%, 90%, and 95%). Next, the samples were placed in Hitachi HCP for critical point drying, coated with platinum, and inspected using a filed emission scanning electron microscope (Hitachi SU-8010).

### Confocal microscopy and living image

For live imaging of sepal development, the main inflorescence was excised from six-to eight-week-old plants containing the *pPRS::GFP-GUS, pAS2*^*WT*^::*GFP-GUS* and *pAS2*^*as2-7D*^::*GFP-GUS* markers (*pPRS::GFP-GUS* in WT and *as2-7D, pAS2*^*WT*^::*GFP-GUS* and *pAS2*^*as2-7D*^::*GFP-GUS* in WT). Siliques were removed with surgical scissors. Flowers above stage 8 were dissected off using tweezers under a stereo microscope. The dissected inflorescences were inserted upright into plates containing live imaging medium (1/2 MS medium supplemented with a 1000-fold dilution of a plant preservative mixture). The plates were filled with autoclaved water and left for one to two hours to rehydrate the dissected inflorescences. Residual buds were further trimmed to early stages with a needle in the water. Recover the inflorescences from dissection by transferring them to fresh media plates for at least half an hour before live imaging. After recovery, the dissected tissue used for live imaging was positioned at an angle to ensure that the flower of the selected stage was directly facing the objective of the confocal microscope. Whole flowers were immersed in autoclaved water containing a 1000-fold diluted plant preservative mixture and imaged using a vertical confocal microscope with a 40x water dipping objective. The following settings were used: excitation laser at 488 nm, emission collection range of 500-550 nm, and a z-step of 0.7-1.0 µm. All confocal imaging were performed with a Nikon C2si confocal. Images were processed by ImageJ (https://imagej.net/software/imagej/) and MorphographX (de Reuille *et al*., 2015).

### Vector construction and plant transformation

*pPRS::GFP-GUS* was constructed by inserting the *PRS* promoter sequence into the binary vector *p35S::GUS-GFP* (Chang et al., 2024) using the *Hind*III and *Sal*I cutting sites. To generate *p35S::PRS*, the *PRS* coding sequence from the start codon to the stop codon was amplified from WT genomic DNA. The PCR product was cloned into *pCAMBIA1300-35S* (a binary vector derived from pCAMBIA1300, containing the *2× CaMV 35S* promoter and the *CaMV* terminator) between the *Kpn*I and *Pst*I cutting sites. The 35S-PRS fragment from the resultant vector was then cut with *Hind*III and *EcoR*I and ligated into the binary vector pCAMBIA2300 (CAMBIA). *pAS2*^*WT*^::*GFP-GUS* and *pAS2*^*as2-7D*^::*GFP-GUS* have been described previously (Yadav *et al*., 2024). All primers used for were listed in Supplementary Table S1.

### RNA extraction and reverse transcription quantitative PCR (RT-qPCR)

Total RNA was extracted from inflorescences (with mature flowers removed) using the Easy Plant RNA Kit (Easy-Do, CAT DR0406050) according to the manufacturer’s instructions. First-strand of cDNA was synthesized using HiScript® II Reverse Transcriptase (Vazyme, CAT R333-01) and then used as a template for RT-qPCR. qPCR was performed in a Bio-Rad CFX96 Real-Time PCR System (Bio-Rad) using Hieff qPCR SYBR Green Master Mix (Yeasen, CAT 11201ES08) according to the manufacturer’s instructions. Quantification of the UBQ gene was used as a control. All primers used for qRT-PCR are listed in Supplementary Table S1.

### Yeast one-hybrid assay

To make the yeast one-hybrid assay bait constructs, the *PRS* promoter sequence were amplified from genomic DNA using specific primers (Supplementary Table S1). The PCR products were ligated into the *pHIS2* vectorusing *Sac*I and *EcoR*I cutting sites to construct the *pPRS::HIS2* (PRS-HIS2). The *AS2* coding sequence (CDS) was inserted into *pGADT7* vector (AD) using *Xho*I and *EcoR*I cutting sites to construct the AS2-AD. The combinations of PRS-HIS2 and AS2-AD, as well as PRS-HIS2 and AD were transformed into yeast strain Y187. Yeast synthetic drop-out media (SD) lacking leucine (Leu) and tryptophan (Trp; Coolaber, PM2222) were used to identify transformed colonies. Three days after transformation, selective colonies were grown in -Leu and -Trp liquid media for 24 hours. The cell suspension was then pipetted onto yeast SD media lacking histidine (His), Leu, and Trp (Coolaber, PM2222) supplemented with 80 mM 3-amino-1,2,4-triazole (3-AT).

### Yeast two-hybrid assay

To confirm the interaction between AS2 and PRS, the CDS of *PRS* gene was amplified and cloned into the *pGBKT7* vector (BD) using *Pst*I and *EcoR*I to construct the PRS-BD and the CDS of *AS2* gene was inserted into *pGADT7* vector using *Xho*I and *EcoR*I to construct the AS2-AD. The PCR primers used for plasmid construction are listed in Supplementary Table S1. The combinations of PRS-BD and AS2-AD, PRS-BD and AD, BD and AS2-AD, and AD and BD were transformed into yeast strain AH109. Yeast SD media lacking Leu and Trp (Coolaber, PM2222) was used to identify transformed colonies. Three days after transformation, selective colonies were grown in -Leu and -Trp liquid media for 24 hours. Then, cell suspension was pipetted onto yeast SD media lacking His, adenine (Ade), Leu, and Trp (Coolaber, PM2222).

### Dual-luciferase assay

The dual-luciferase assays were performed in *Nicotiana benthamiana* (*N. benthamiana*) leaves as previously described (Yin *et al*., 2010). To generate the constructs for dual-luciferase assays, the promoter region of *PRS* was cloned into the *pGreen*II *0800-LUC* vector using *BamH*I and *Hind*III to construct the *pGreen*II *0800-PRSpro-LUC* (PRS-LUC), and the CDS of *AS2* was cloned into the *pGreenII 0029-62-SK* (SK) (Yin *et al*., 2010) vector using *BamH*I and *Hind*III to construct *pGreenII 0029-62-AS2-SK* (AS2-SK). Both constructs were transferred into *Agrobacterium tumefaciens* strain GV3101 (pSoup) and used to infiltrate *N. benthamiana* leaves. Infiltrated plants were incubated at 22□ for 72 hours. For qualitative observations, the LUC images were captured using a low-light cooled CCD imaging apparatus (Tanon-5200 with AllDoc_x software). Transformed leaves were sprayed with 1 mM luciferin and placed in darkness for several minutes before luminescence detection. For quantitative analysis, firefly and renilla luciferase activities were monitored using the Dual-Luciferase Reporter Assay System (Promega, E1910) on the dual fluorescence detector (GloMax96).

### Firefly luciferase complementation imaging assay

The transient assays were performed in *N. benthamiana* leaves as previously described (Zhao *et al*., 2019). The CDS of *AS2* gene was PCR-amplified and cloned into the *pCambia1300-35S-nLuc* vector using *Sac*I and *Sal*I to construct the AS2-nLUC, and the CDS of *PRS* gene was PCR-amplified and cloned into the *pCambia1300-35S-cLuc* vector using *Sac*I and *Sal*I to construct the PRS-cLUC. Both constructs were transferred into *Agrobacterium tumefaciens* strain GV3101 and used to infiltrate *N. benthamiana* leaves. Infiltrated plants were incubated at 22°C for 72 hours. The transformed leaves were sprayed with 1 mM luciferin and placed in darkness for several minutes before luminescence detection. The LUC images were captured using a low-light cooled CCD imaging apparatus (Tanon-5200 with AllDoc_x software).

### BiFC assay

The BiFC assays were performed in *N. benthamiana* leaves as previously described (Yang *et al*., 2007). To generate the constructs for BiFC assays, the coding regions of *PRS* and *AS2* were cloned in frame into the *BamH*I and *Sal*I digested *p2YN* and *p2YC* (Yang *et al*., 2007) vectors to generate PRS-cYFP and nYFP-AS2, respectively. Both constructs were transferred into *Agrobacterium tumefaciens* strain GV3101 and used to infiltrate *N. benthamiana* leaves. The fluorescence signals were examined by using a Nikon C2si confocal.

## RESULTS

### *PRS* gene deletion suppresses outgrowth formation on *as2-7D* sepal abaxial epidermis

To investigate the molecular mechanism involved in maintaining sepal flatness in *Arabidopsis*, we performed a suppressor screen on *as2-7D* and isolated a suppressor mutant of *as2-7D* termed *as2-7D ssp1*. Compared to *as2-7D* sepals, *as2-7D ssp1* sepals are smoother on the outer (abaxial) epidermis, with significantly fewer outgrowths. Compared to WT sepals, *as2-7D ssp1* sepals are narrower, exposing a part of the internal floral organs (Fig. 1A to 1C). Using the QTL-seq mapping strategy, the candidate region was narrowed down to 9-16 Mb on chromosome 2 (Supplementary Fig. S1). In this region, a large segment deletion of 25,189 bp closely linked to the *ssp1* mutation was identified in the *as2-7D ssp1* genome (Fig. 1D). This deletion disrupted the coding regions of six genes, among which only the *PRS* (*At2G28610*) gene has been reported to be involved in sepal morphogenesis (Matsumoto and Okada, 2001). Consequently, *PRS* was selected as the candidate gene of *SSP1*. Since two *prs* mutants have been previously reported, the *as2-7D ssp1* mutant was renamed as *as2-7D prs-3*. The *prs-3* single mutant was isolated in the F_2_ generation by crossing *as2-7D prs-3* with WT. Sepals of *prs-3* mutants have a smooth epidermis and are narrower compared to WT sepals (Fig. 1A to 1C; Supplementary Fig. S2), consistent with the phenotype of other *prs* alleles (Nakata *et al*., 2012). To further verify that the *PRS* gene is indeed the *SSP1* gene, we crossed *as2-5D* (an *AS2* mutant in the *Col* background with the same point mutation as *as2-7D*) with *prs-4* (a *prs* mutant in the *Col* background) and isolated the *as2-5D prs-4* double mutant. Phenotypic observations of *Col-0, as2-5D, prs-4*, and *as2-5D prs-4* flowers revealed that *Col-0* and *prs-4* sepals had smooth surfaces. In contrast, *as2-5D* sepals formed outgrowths on the surface, though fewer than those on *as2-7D* sepals, while *as2-5D prs-4* sepals were smooth and flat without obvious outgrowth formation (Figure. 1E). These results indicate that *PRS* mutation indeed suppresses the outgrowth formation on the abaxial sepal epidermis caused by ectopic *AS2* expression.

**Figure 1.**
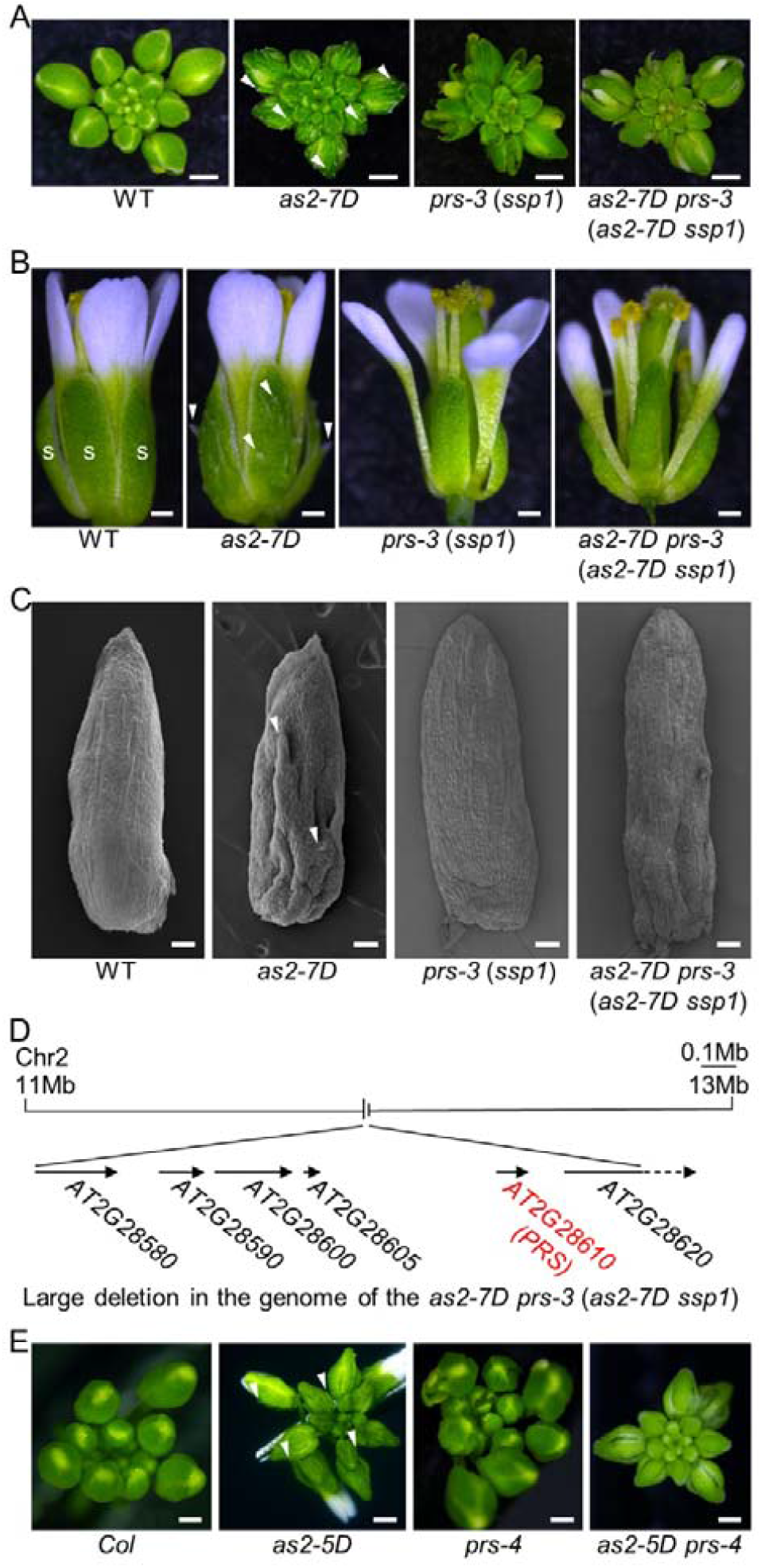
*PRS* gene deletion suppresses outgrowth formation on *as2-7D* sepal abaxial epidermis. (A) Inflorescences of WT, *as2-7D, prs-3* (*ssp1*) and *as2-7D prs-3* (*as2-7D ssp1*). *as2-7D* sepals have many outgrowths on the surface; *prs-3* (*ssp1*) sepals are narrower compared with WT sepals; the *as2-7D prs-3* (*as2-7D ssp1*) sepals are smoother on the abaxial epidermis compared to *as2-7D* sepals. Scale bars: 400 μm. (B) Flowers of WT, *as2-7D, prs-3* (*ssp1*), and *as2-7D prs-3* (*as2-7D ssp1*). Sepals (green leaf-like organs) are marked with the letter “S” in white in WT. Scale bars: 200 μm. (C) Scanning electron microscope images of sepals of WT, *as2-7D, prs-3* (*ssp1*) and *as2-7D prs-3* (*as2-7D ssp1*). Scale bars: 100 μm. Arrowheads in A-C point to the outgrowths on *as2-7D* sepals. (D) The large fragment deletion in the genome of the *as2-7D prs-3* (*as2-7D ssp1*) includes six genes, among which *AT2G28610* (*PRS*) is the gene involved in the suppression of the *as2-7D* sepal phenotype. (E) Inflorescences of *Col, as2-5D, prs-4*, and *as2-5D prs-4. as2-5D* sepals have outgrowths on the surface (as arrowheads point at). Scale bars: 400 μm.

### *AS2* up-regulates *PRS* expression on *as2-7D* sepal abaxial epidermis

The abaxial epidermal outgrowths in *as2-7D* sepals are caused by ectopic *AS2* expression on the abaxial epidermis. To investigate how the *PRS* deletion suppresses outgrowth formation on *as2-7D* sepal abaxial epidermis, we first examined whether *PRS* deletion affected *AS2* expression pattern. We drove green fluorescent protein (GFP) expression under the WT *AS2* promoter (*pAS2*^*WT*^::*GFP-GUS*) and the mutant *as2-7D AS2* promoter (*pAS2*^*as2-7D*^::*GFP-GUS*) in *prs-3*, and obtained the *pAS2*^*WT*^::*GFP-GUS prs-3* and *pAS2*^*as2-7D*^::*GFP-GUS prs-3* reporter lines. Confocal microscopy showed that the WT *AS2* promoter drives *GFP* expression specifically on the adaxial sepal epidermis of *prs-3* (Fig. 2A,2C, and 2E), while the mutant *AS2* promoter drives *GFP* expression on both the adaxial and abaxial epidermal layers of *prs-3* (Fig. 2B,2D, and 2F). The expression patterns of *pAS2*^*WT*^::*GFP-GUS* and *pAS2*^*as2-7D*^::*GFP-GUS* in *prs-3* sepals are similar to their expression patterns in WT sepals (Yadav *et al*., 2024), indicating that the *prs-3* mutation does not alter the activity of WT and mutant *AS2* promoters. Therefore, the suppression of outgrowth formation on the abaxial epidermis of *as2-7D prs-3* sepals by *prs-3* mutation is not achieved by suppressing the ectopic expression of *AS2* on sepal abaxial epidermis. Instead, *PRS* is required for the ectopically expressed *AS2* to induce outgrowths on the sepal abaxial epidermis.

**Figure 2.**
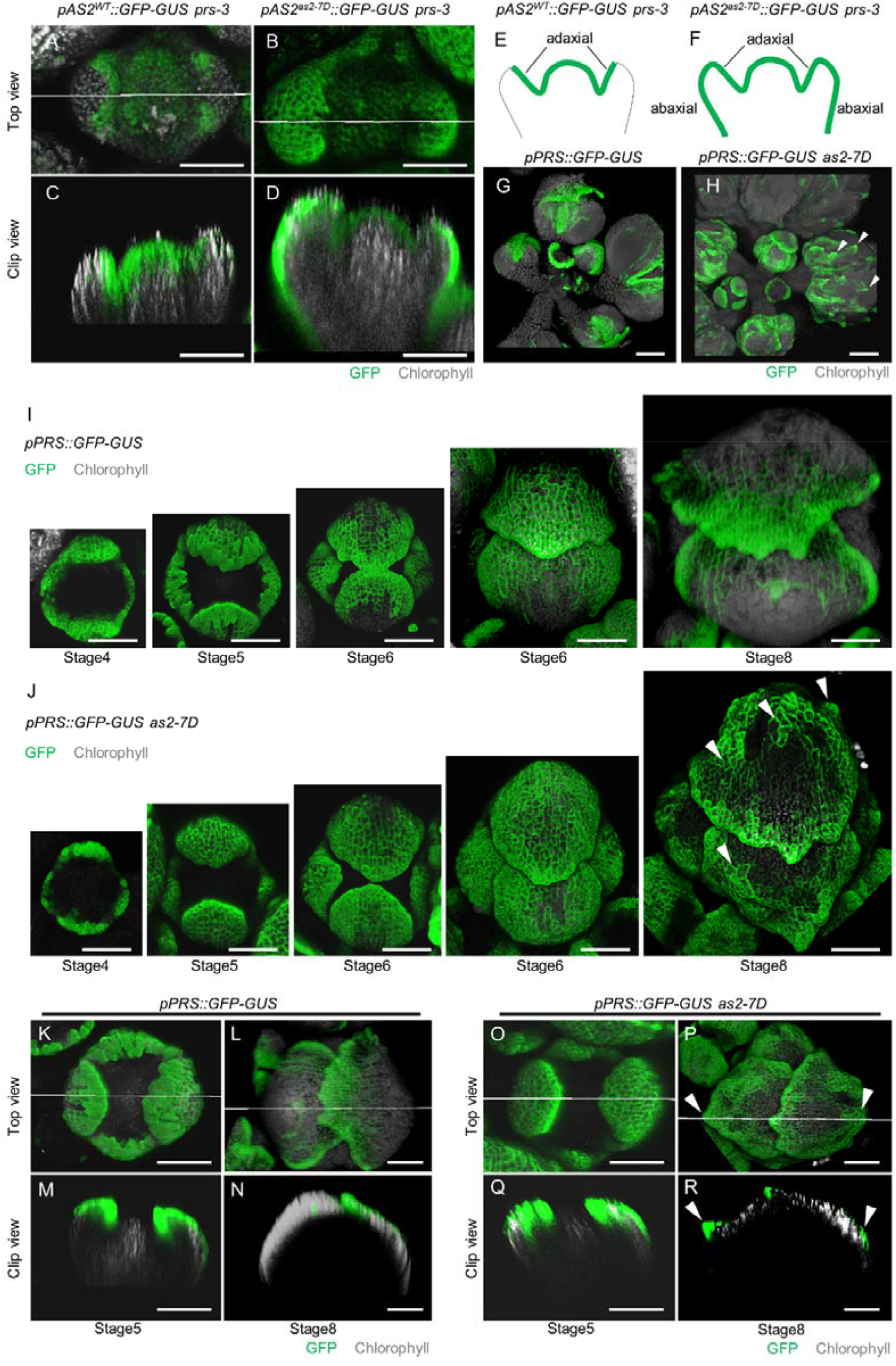
*AS2* up-regulates *PRS* expression on *as2-7D* sepal abaxial epidermis. (A to F) Confocal images showing the WT *AS2* promoter (*pAS2*^*WT*^::*GFP-GUS*) and the mutant AS2 promoter (*pAS2*^*as2-7D*^::*GFP-GUS*) activity in *prs-3* flowers. The WT *AS2* promoter drives GFP expression on the adaxial sepal epidermis of *prs-3* (A, C, E). The mutant *as2-7D* promoter drives GFP expression on both the adaxial and abaxial epidermal layers of *prs-3* (B, D, F). A and B are top views; C and D are the side views of clips at the white lines in the top views; E and F are sketches of GFP distribution patterns. GFP: green; Chlorophyll: gray. Scale bars: 400 μm. (G and H) Confocal images showing the *PRS* promoter (*pPRS::GFP-GUS*) activity in WT (G) and *as2-7D* (H) inflorescences. *PRS* is expressed at the margins of WT sepals, whereas at the margins and outgrowths (as arrowheads show in H) of *as2-7D* sepals. GFP: green; Chlorophyll: gray. Scale bars: 100 μm. (I and J) Confocal images of the *PRS* promoter (*pPRS::GFP-GUS*) activity in WT and *as2-7D* sepals at different developmental stages. *PRS* is widely expressed in all four sepals at stages 4 and 5 in both WT and *as2-7D* sepals. In WT sepals at stage 6 and later, *PRS* expression is restricted to the sepal margin, whereas *PRS* is expressed at both the margins and middle regions in *as2-7D* sepals, especially at outgrowths (as arrowheads point). GFP: green; Chlorophyll: gray. Scale bars: 50 μm. (K to R) Longitudinal sections of WT and *as2-7D* flowers at stage 5 and stage 8. *PRS* is widely expressed in WT (K and M) and *as2-7D* sepals at stage 5 (O and Q). In WT sepals at stage 8, *PRS* is expressed at the margins of the sepals (L and N). In *as2-7D* sepals at stage 8, *PRS* is expressed at the margins and the outgrowths (as arrowheads point at) (P and R). K and L are reproduced from I. O and P are reproduced from J. M, N, Q, and R are the side views of clips at the while lines in K, L, O, and P respectively. GFP: green; Chlorophyll: gray. Scale bars: 50 μm.

Our results suggest that *PRS* functions downstream of *AS2* to influence sepal flatness. To further explore this hypothesis, we examined whether *PRS* expression was affected by the *as2-7D* mutation. Using RT-qPCR, we quantified the expression levels of *PRS* in WT and *as2-7D* sepals. The results showed that *PRS* expression level was significantly higher in *as2-7D* sepals than in WT sepals (Supplementary Fig. S3). *PRS* is a member of the WOX transcription factor family, whose genes exhibit specific expression patterns critical to their functional divergence, and previous study has shown that *PRS* displays a dynamic expression pattern throughout sepal development using RNA *in situ* hybridization (Matsumoto and Okada, 2001). To explore how ectopic *AS2* expression up-regulates *PRS*, we generated *PRS* transcriptional reporter lines to compare *PRS* expression patterns in WT (*pPRS::GFP-GUS*) and *as2-7D* (*pPRS::GFP-GUS as2-7D*) sepals. Confocal microscopy showed that overall *PRS* was expressed at the margins of WT sepals (Fig. 2G) and at both the margins and outgrowths of *as2-7D* sepals (Fig. 2H). Observations of individual flowers at different developmental stages showed similar *PRS* expression patterns in WT and *as2-7D* sepals at stages 4 and 5, with broad expression in all four sepals (Fig. 2I, and 2J). However, starting from stage 6, *as2-7D* sepals exhibited divergent *PRS* expression patterns compared to WT sepals. In WT sepals at stage 6 and beyond, *PRS* expression was restricted to the sepal margins (Fig. 2I), consistent with the expression patterns observed in previous research through RNA *in situ* hybridization (Matsumoto and Okada, 2001). In contrast, *as2-7D* sepals at similar stages displayed broader *PRS* expression, with expression at both the margins and the outgrowths (Fig. 2J). Furthermore, longitudinal sections of sepals from stage 5 and stage 8 showed that *PRS* was widely expressed on both the adaxial and abaxial sides of sepals in WT and *as2-7D* at stage 5 (Fig. 2K,2M,2O, and 2Q). At stage 8, *PRS* expression was restricted at the sepal margins in WT (Fig. 2L and 2N), whereas *as2-7D* sepals exhibited prominent *PRS* expression at the margins and the outgrowths (Fig. 2P and 2R).

*as2-7D* sepals start to exhibit ectopic *PRS* expression at stage 6, which coincides with the developmental stage when the abaxial epidermis of *as2-7D* sepals starts to form outgrowths. This timing coincidence suggests a potential relationship between the altered *PRS* expression pattern and epidermal outgrowth formation. To investigate the connection between ectopic *PRS* expression and outgrowth formation on *as2-7D* sepals, we tracked *PRS* expression patterns before and after the outgrowth formation. Our observations revealed that ectopic *PRS* expression consistently preceded outgrowth formation. Before the formation of outgrowths (prior to stage 6), *PRS* was expressed throughout the sepal abaxial epidermis. As *as2-7D* sepals continued to develop and outgrowths began to form on the abaxial epidermis, *PRS* expression activity gradually receded toward the sepal margins but remained at the outgrowths (Fig. 3A and 3B).

**Figure 3.**
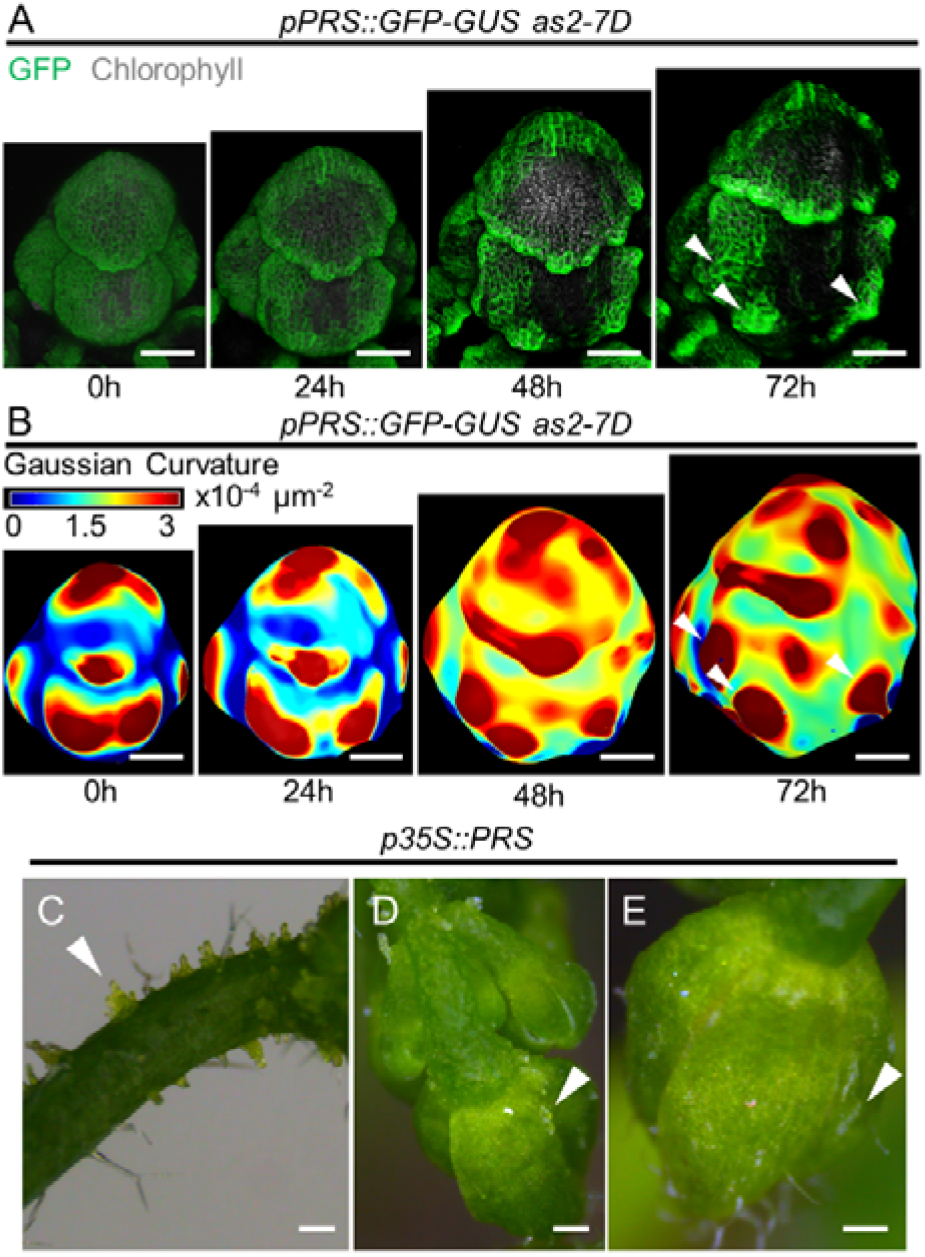
*PRS* promotes outgrowth formation on the sepal epidermis. (A) Confocal images of the same flower from *pPRS::GFP-GUS as2-7D* at 0, 24, 48, and 72 hours, respectively. GFP: green; Chlorophyll: gray. Note the *GFP* expression at the outgrowths (arrowheads). (B) Gaussian curvature of the flowers in A. Warm colors denote out-of-plane (bulging) and cool colors represent inward (saddling) deformations. At 0 hour, *PRS* is widely expressed on the abaxial epidermis of sepals, and the sepal exhibits relatively uniform surface curvature. At 24 and 48 hours, *PRS* expression gradually retreats toward the sepal margin, and outgrowths initiate on the sepal abaxial epidermis. At 72 hour, as *PRS* expression retreats further to the margins, it lingers on the rising outgrowths (as arrowheads show). Scale bars:50 μm. (C to E) Phenotypes of *p35S::PRS* transgenic plants. (C) Outgrowths (arrowhead) on the peduncle. (D) Outgrowths (arrowhead) on the sepal. (E) White wrinkle structures (arrowhead) on the sepal. Scale bars: 400 μm.

In conclusion, our tracking of *PRS* expression during early sepal development supports a model in which *PRS* expression activity gradually recedes toward the sepal margins as sepals grow in WT, while in *as2-7D* sepals the ectopically expressed AS2 on the abaxial epidermis tends to detain *PRS* expression and keeps it to linger on the rising outgrowth. So we believe that AS2 up-regulates *PRS* expression on *as2-7D* sepal abaxial epidermis,

### *PRS* promotes the formation of outgrowths on the sepal epidermis

To further investigate the relationship between up-regulated *PRS* expression and epidermal outgrowth formation, we generated transgenic lines that overexpressed *PRS* gene (*p35S::PRS*) in WT plants. In these transgenic plants, outgrowths were observed on the epidermis of the peduncles and sepals (Fig. 3C to 3E), showing that increased *PRS* expression can lead to outgrowth formation on sepals. Combined with the *PRS* expression pattern analysis, this result provides evidence that the ectopic *PRS* expression causes epidermal outgrowth formation in *as2-7D* sepals. Introducing *p35S::PRS* into *as2-101*, a loss-of-function mutant of *AS2*, also resulted in outgrowth formation on the sepal epidermis (Supplementary Fig. S4), indicating that *PRS* promotes epidermal outgrowth formation independently of functional *AS2*. This finding supports our previous conclusion that *PRS* functions downstream of *AS2* to influence sepal flatness.

### AS2 directly promotes *PRS* expression

Expression analysis demonstrated that ectopic *AS2* expression up-regulates *PRS* in *as2-7D* sepals. A yeast one hybrid assay was performed to investigate whether *AS2* up-regulates *PRS* directly. Results showed that AS2 can bind to the promoter region of *PRS* (Fig. 4A). Dual-luciferase assays in *N. benthamiana* leaves were further performed to test the effects of AS2 on the transcription of PRS-LUC reporter, which displayed that AS2 promoted the expression of *PRS* (Fig. 4B and 4C).

**Figure 4.**
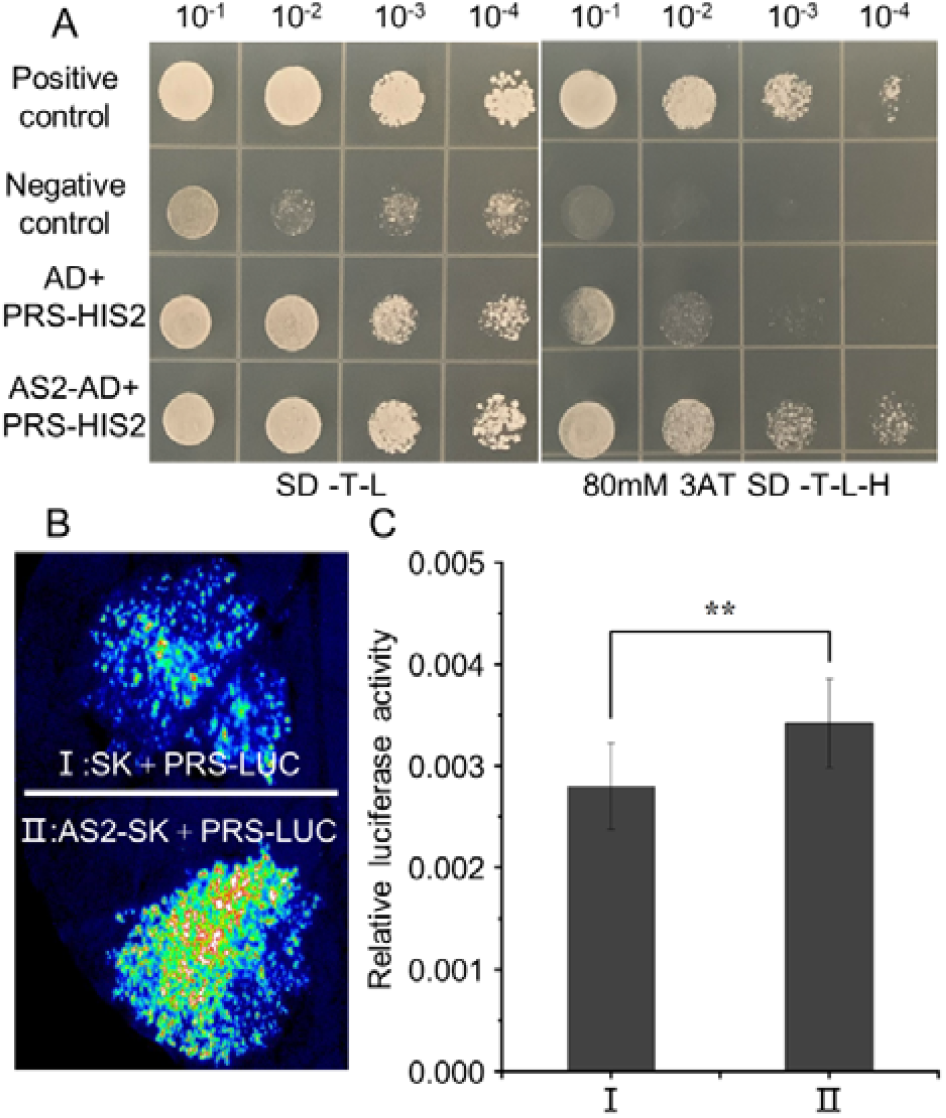
AS2 promotes *PRS* expression directly. (A) AS2 binds to the *PRS* promoter region in the yeast one-hybrid assay. Transformed yeast cells were spotted on SD -Trp-Leu or SD -Trp-Leu-His medium supplemented with 80 mM 3AT in 10-, 100-, 1000- and 10000-fold dilutions. The pGADT7-T and pGBKT7-53 were used as positive controls, while the pGADT7-T and pGBKT7-lam vectors served as negative controls. (B) Transient expression assay showing that AS2 activates *PRS*. Representative image of a *N. benthamiana* leaf 72 hours after infiltration was shown. (C) Quantitative analysis of luminescence intensity in (B). Eleven independent replicates were performed. Error bars represent standard deviation (SD). Data are presented as mean ± SD for more than three independent experiments. Two-tailed Student’s t-test, \*\**p* <0.01.

Taken together, these results indicate that AS2 promotes *PRS* expression through direct binding to *PRS* promoter regions.

### AS2 physically interacts with PRS

Previous studies have reported that AS2 physically interacts with transcription factor TEOSINTE BRANCHED 1, CYCLOIDEA, AND PCF FAMILY 4 (TCP4) (Li *et al*., 2012) and that PRS also interacts with TCP4 physically (Wanamaker *et al*., 2017). Therefore, we explored whether AS2 could interact with PRS at the protein level. In the yeast two-hybrid system, yeast AH109 colonies co-transformed with AS2-AD and PRS-BD grew well on selective medium (Fig. 5A), indicating that AS2 interacts directly with PRS in yeast. Luciferase complementation imaging assays further showed that AS2-nLUC interacted with PRS-cLUC (Fig.5B). Bimolecular fluorescence complementation (BiFC) assays displayed that YFP fluorescence was detected in the nuclei of *N. benthamiana* leaves co-transformed with AS2-nYFP and PRS-cYFP, but not in those of tobacco leaves transformed with the negative controls (Fig. 5C), further confirming that AS2 can physically interact with PRS in the nuclei.

**Figure 5.**
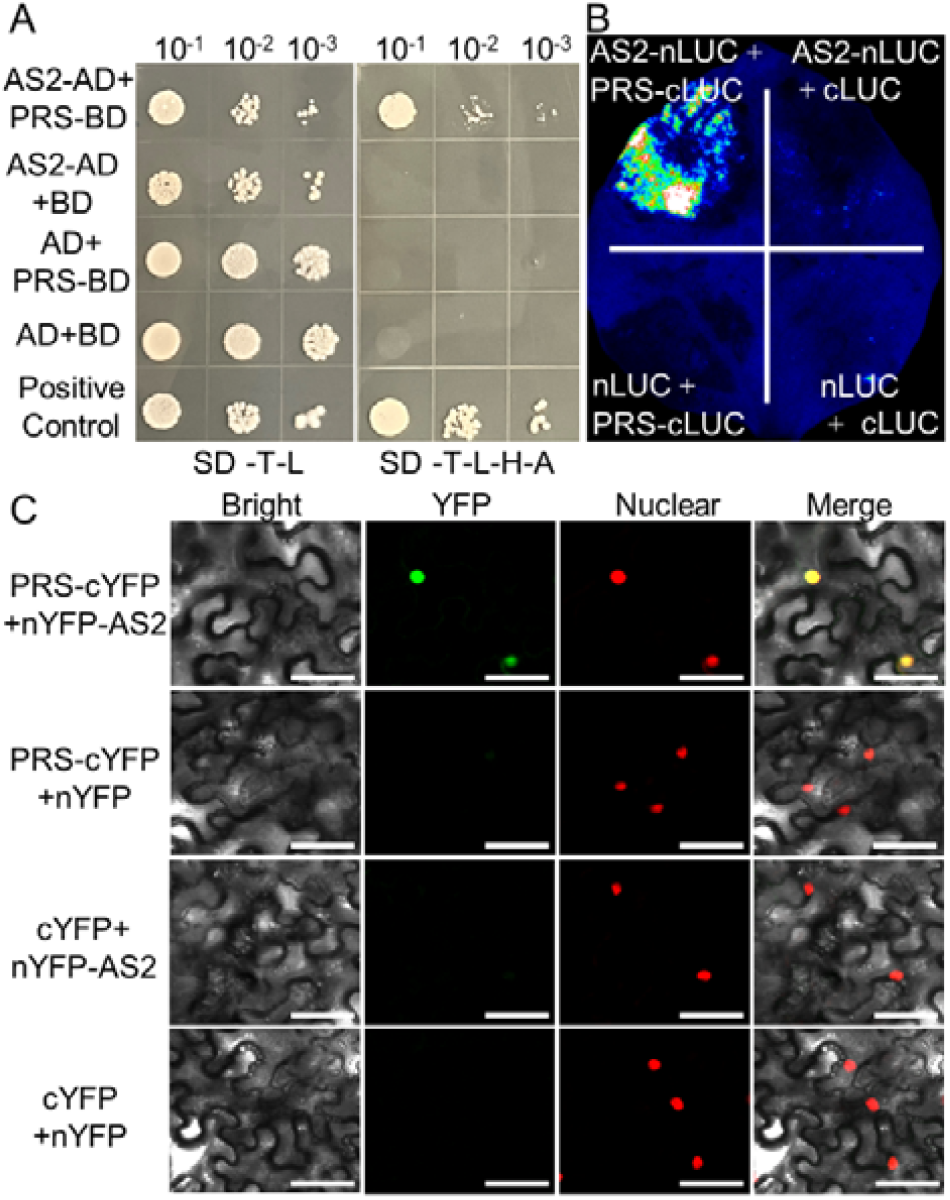
AS2 physically interacts with PRS. (A)AS2 interacted with PRS in the yeast two-hybrid assay. Transformed yeast cells were spotted on SD-Leu-Trp or SD-Leu-Trp-His-Ade medium in 10-, 100-, and 1000-fold dilutions. The pGBKT7-53 and pGADT7-T vector was used as a positive control. (B) Protein-protein interactions between AS2 and PRS were analyzed by dual-Luciferase reporter assay in *N. benthamiana* leaf. (C) BiFC assays in *N. benthamiana* leaves showing the interaction of PRS and AS2 in living cells. The nucleus was indicated by carrying a nuclear localization signal (H2B-RFP). Scale bars: 50 μm.

## DISCUSSION

In this study, through suppressor screening, we demonstrated that loss-of-function of the *PRS* gene suppresses the outgrowth formation on *as2-7D* sepal abaxial epidermis. Using live imaging technology to track the *PRS* expression dynamics and sepal outgrowth formation on *as2-7D* sepal abaxial epidermis, we provided a detailed depiction of the *PRS* expression pattern at a cellular resolution throughout the early developmental stages of sepals, and found that on *as2-7D* sepal abaxial epidermis, *AS2* expression up-regulated *PRS* expression, which resulted in the epidermal outgrowth initiation.

Previous studies have shown that ectopic *AS2* expression disrupts sepal flatness through disturbing the balance of cellular growth and cell mechanics between the two epidermal layers of sepals (Yadav et al., 2024). Our study shows that *PRS*, a WOX family gene, affects sepal flatness downstream of *AS2*. The WOX family genes have been demonstrated to play important regulatory roles in critical developmental stages of plants, such as embryo formation, stem cell stability, and organ formation. Their multifaceted functions in plant development are closely related to their ability to promote cell division or prevent premature differentiation of cells (Deveaux *et al*., 2008; Baesso *et al*., 2020; Zhang *et al*., 2020; Bueno *et al*., 2021). The cellular mechanisms underlying *PRS*’s function in maintaining sepal flatness worth further exploration. Despite active research on the WOX family, detailed molecular mechanisms underlying and gene networks involved in the functions of these transcription factors still need to be deciphered (Lian *et al*., 2014; Tvorogova *et al*., 2021). Our study reveals that *AS2* and *PRS* interact on multiple levels. AS2 and PRS interact physically, and AS2 also activate *PRS* expression through direct binding to *PRS* promoter region. These findings, combined with previous research on the versatile functions of *WOX* genes, suggest that complicated regulatory networks might be involved in role of *AS2* and *PRS* in regulating sepal epidermal flatness. *WOX* family transcription factors, including PRS, contain *WUS-box* motifs that confer them transcriptional repressive activity and may act as transcriptional repressors in plants (Ikeda *et al*., 2009). Does PRS also directly or indirectly target AS2? Does PRS regulate its own transcription by forming a regulatory complex with AS2? What are the downstream genes co-regulated by PRS and AS2 to affect sepal morphology? These are interesting questions that will deepen our understanding of the working mechanisms of *WOX* family proteins and worth to be addressed in the future studies.

Research elucidating the mechanisms of abaxial-adaxial polarity establishment proposes that in *Arabidopsis* leaves, *AS2* is specifically expressed on the adaxial epidermis, while *PRS* is expressed in the middle region (Du *et al*., 2018). However, our research shows that the *AS2* expression region and the *PRS* expression domain overlap at the sepal margin throughout the sepal development. It is reasonable to hypothesize that the interaction between *AS2* and *PRS* on *as2-7D* sepal abaxial epidermis, both on the transcription level and on the protein level, also works in WT sepal margins. However, when the *pPRS::GFP-GUS* vector was transferred into *as2-101* mutant, it was observed that the expression pattern of *PRS* in *as2-101* sepals was the same as that in WT ones (Supplementary Fig. S4), suggesting that the expression of *PRS* at the sepal margin does not require AS2 function. Therefore, probably divergent mechanisms are involved in *PRS* expression regulation in different tissues. It would be interesting to explore whether the interaction between *AS2* and *PRS* also plays a role in sepal margin development.

The *PRS* deletion fails to completely suppress the *as2-7D* phenotype, suggesting that other factors or pathways are involved in modulating sepal epidermal flatness downstream of AS2. In addition, it has been reported that AUXIN RESPONSE FACTOR 3/ETTIN (ARF3/ETT) can suppress *PRS* expression by directly binding to its promoter region (Guan *et al*., 2017), while AS2 modulates the expression level of *ETT/ARF3* by maintaining CpG methylation in specific exons of *ETT/ARF3* (Vial-Pradel *et al*., 2018; Iwakawa *et al*., 2020). Hence, it is highly possible that that *AS2* might also regulate *PRS* expression indirectly, probably through *ARF3*.

## SUPPLEMENTARY DATA

The following supplementary data are available online.

Figure S1. QTL-seq mapping of the *SSP1* gene.

Figure S2. The morphology of WT and *prs-3* sepals.

Figure S3. RT-qPCR analysis of *PRS* expression in WT and *as2-7D* sepals.

Figure S4. The phenotype of *p35S: PRS as2-101* transgenic plant.

Table S1. Primers used in this study.

### ACKNOWLEDGMENTS

We thank Prof. Juan Xu for providing the confocal microscope, Prof. Songlin Bai for providing the *pGreenII 0800-Luc* and *pGreenII 0029-62-SK* plasmids, Prof. Xiaobo Zhao for providing the *pCambia1300-35S-nLuc* and *pCambia1300-35S-cLuc* plasmids, Prof. Dianxing Wu, Prof. Xiaoli Shu and Prof. Jingsong Bao for providing experimental facilities, Prof. Ming Zhou for the comments on the article.

## AUTHOR CONTRIBUTIONS

**Ruoyu Liu:** Conceptualization, Methodology, Formal analysis, Investigation, Resources, Data curation, Writing – review & editing, Writing – original draft, Visualization, Validation. **Zeming Wang:** Conceptualization, Methodology, Formal analysis, Investigation, Resources, Data curation, Writing – Validation. **Xi He:** Methodology, Resources, Investigation. **Heng Zhou:** Investigation. **Yiru Xu:** Resources. **Lilan Hong:** Writing – review & editing, Supervision, Project administration, Funding acquisition, Data curation, Conceptualization.

