## Supplemental Table S1 for "*PRESSED FLOWER* works downstream of *ASYMMETRIC LEAVES 2* to affect sepal flatness in *Arabidopsis*"

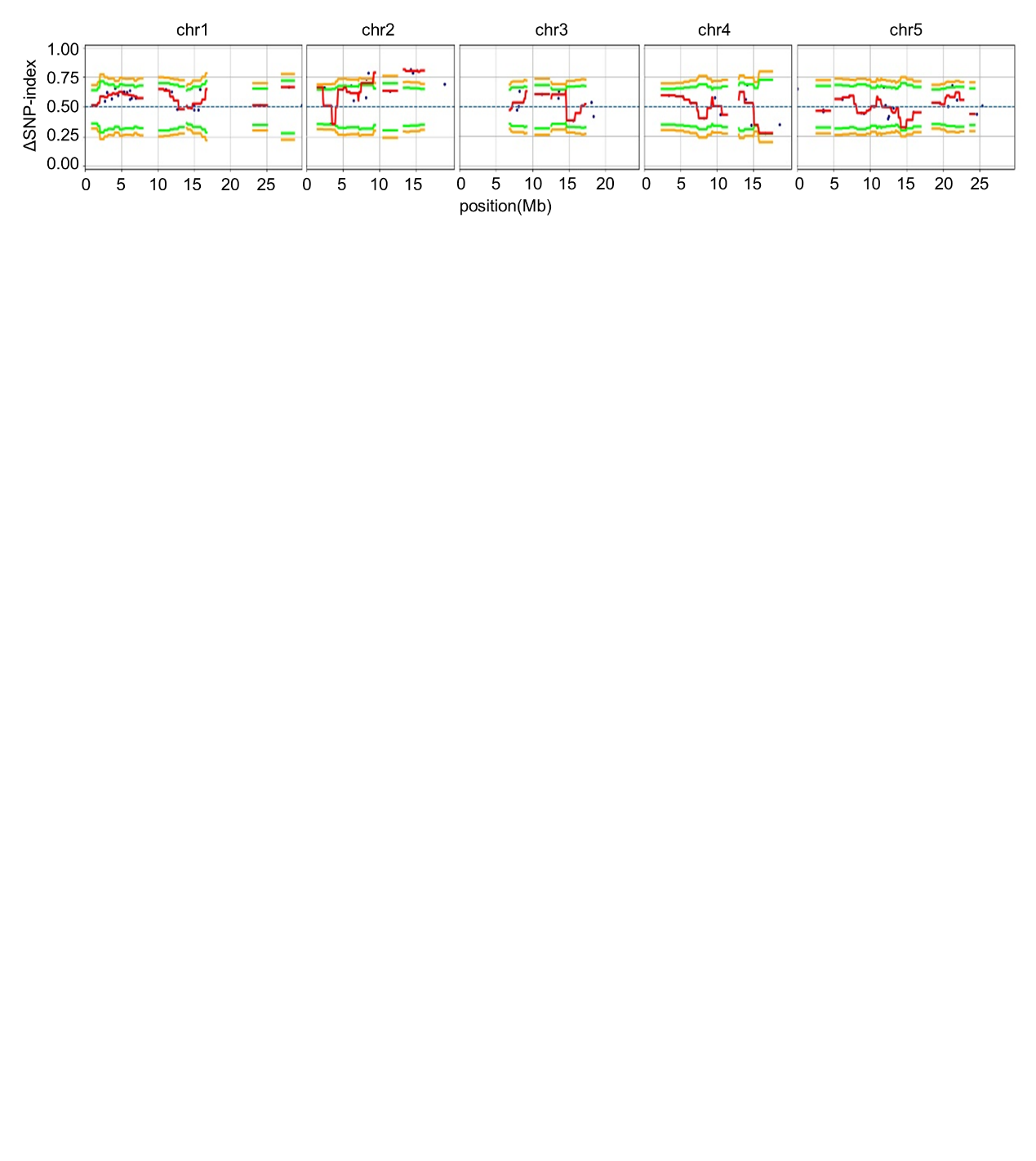

**Figure S1.** **QTL-seq mapping of the *SSP1* gene**

Results obtained by the QTL-seq method. ΔSNP index plots of all the five chromosomes with statistical confidence interval under the null hypothesis of no QTLs (orange, *p* < 0.01; and green, *p*< 0.05).

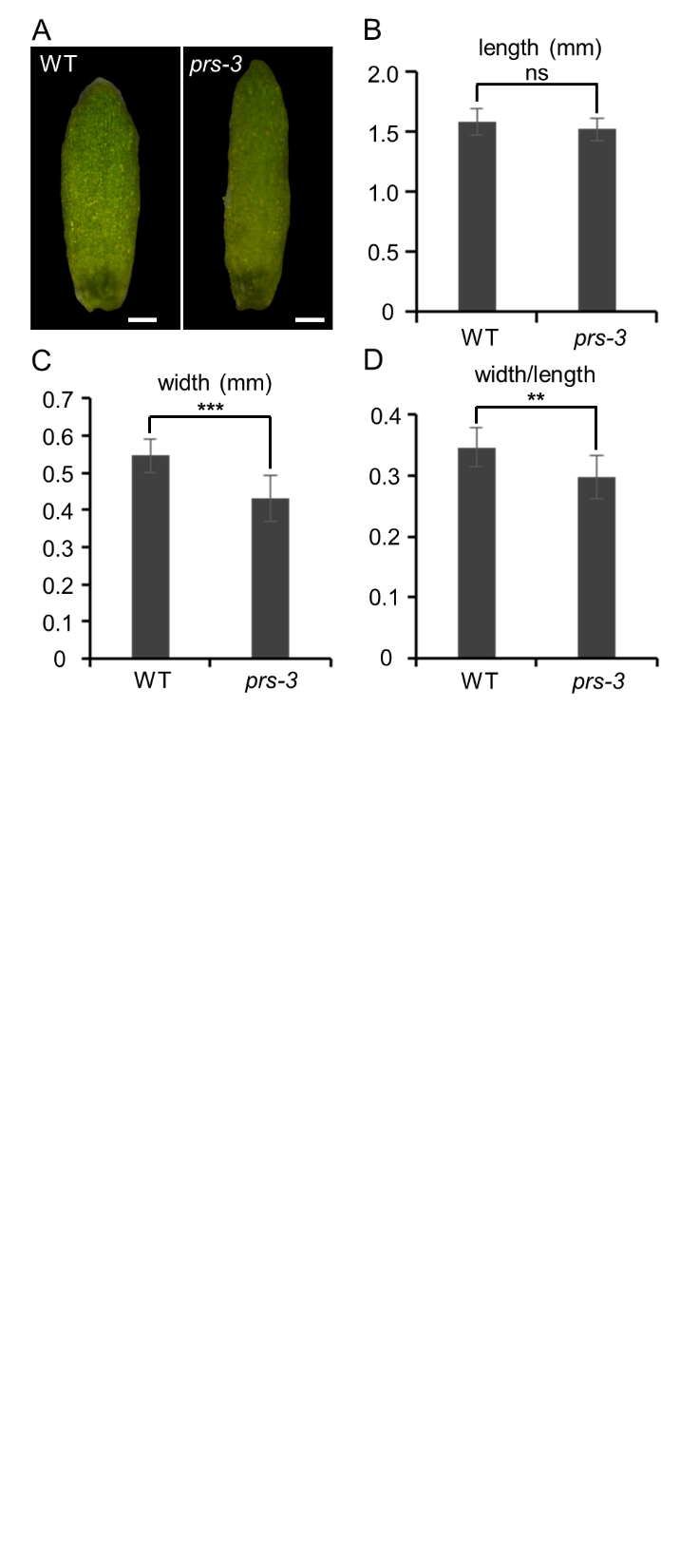

**Figure S2. The morphology of WT and *prs-3* sepals**

(A) The morphology of the WT and *prs-3* abaxial sepals. Scale bars: 200 μm. (B) There is no significant difference in sepal length between WT and the *prs-3* mutant. (C) The *prs-3* sepal is narrower than the WT sepal. (D) The *prs-3* sepal has a smaller width-to-length ratio than the WT sepal. In B-D, thirteen WT sepals and twelve sepals *prs-3* were measured. Data are presented as mean ± SD. ns means not significant with *p*＞0.05; ***p* < 0.01; ****p* < 0.001; Two-tailed Student’s t-test.

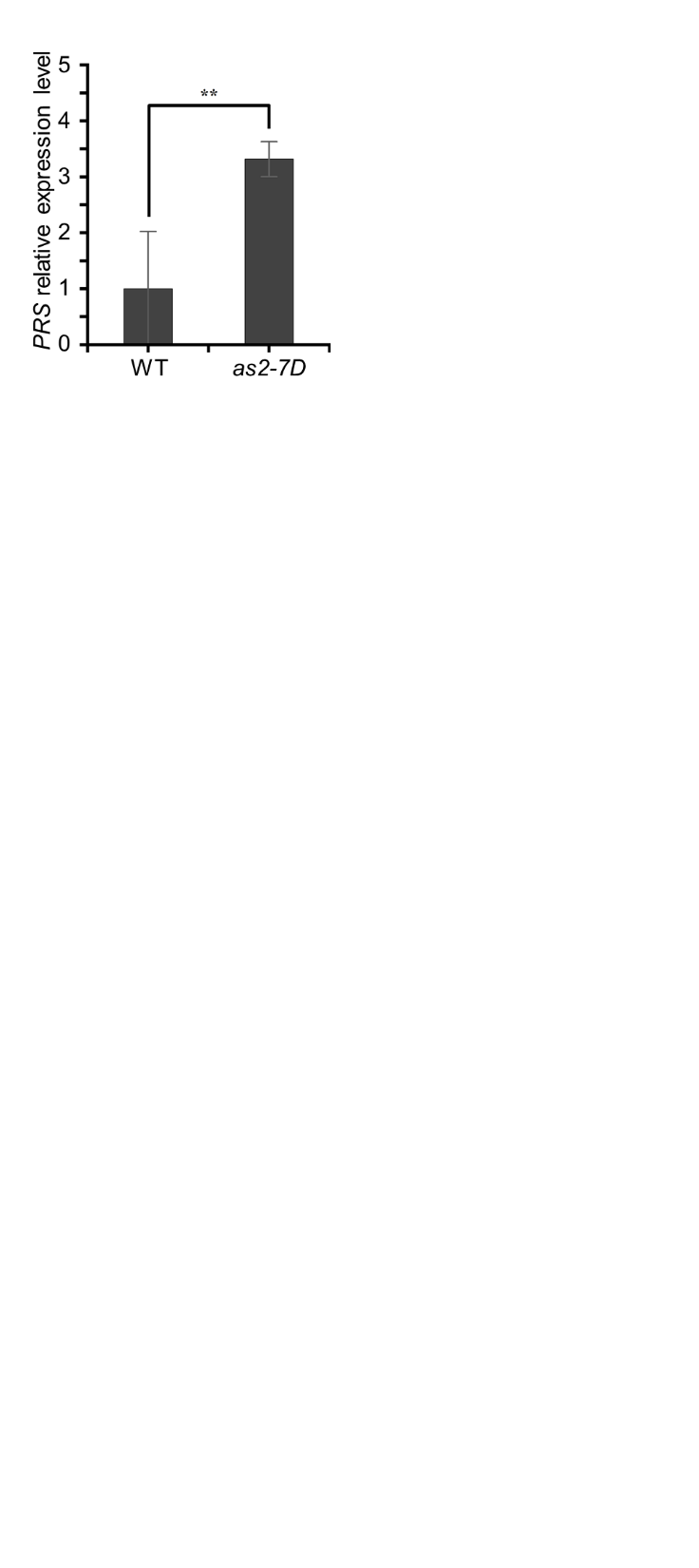

**Figure S3. RT-qPCR analysis of *PRS* expression in WT and *as2-7D* sepals.**

Data are presented as mean ± SD for three independent experiments. Two-tailed Student’s t-test, ***p* < 0.01.

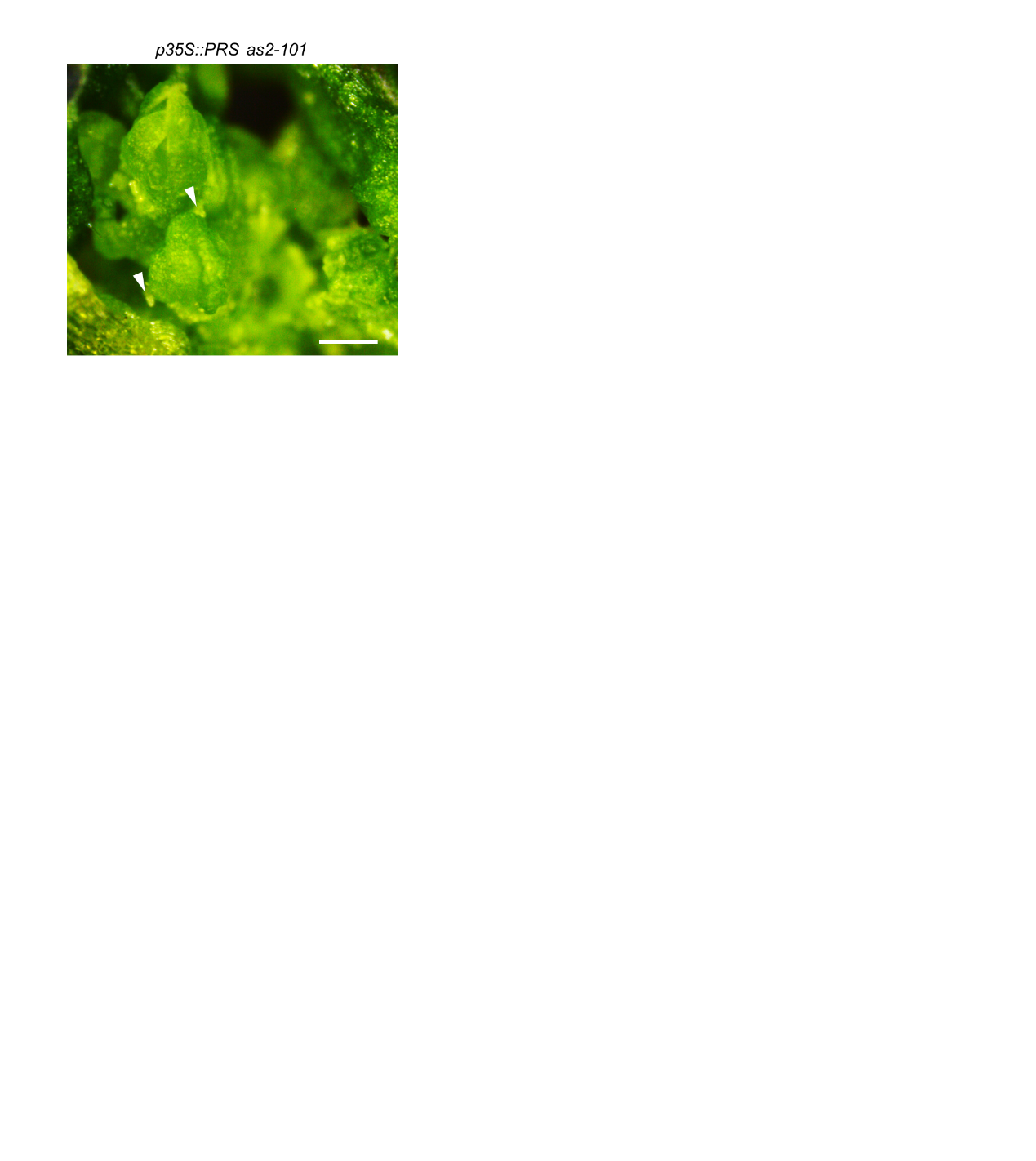

**Figure S4. The phenotype of *p35S: PRS as2-101* transgenic plant**

Multicellular outgrowths on the sepal (white arrowheads). Scale bar: 200 μm.

**Table S1. Primers used in this study**

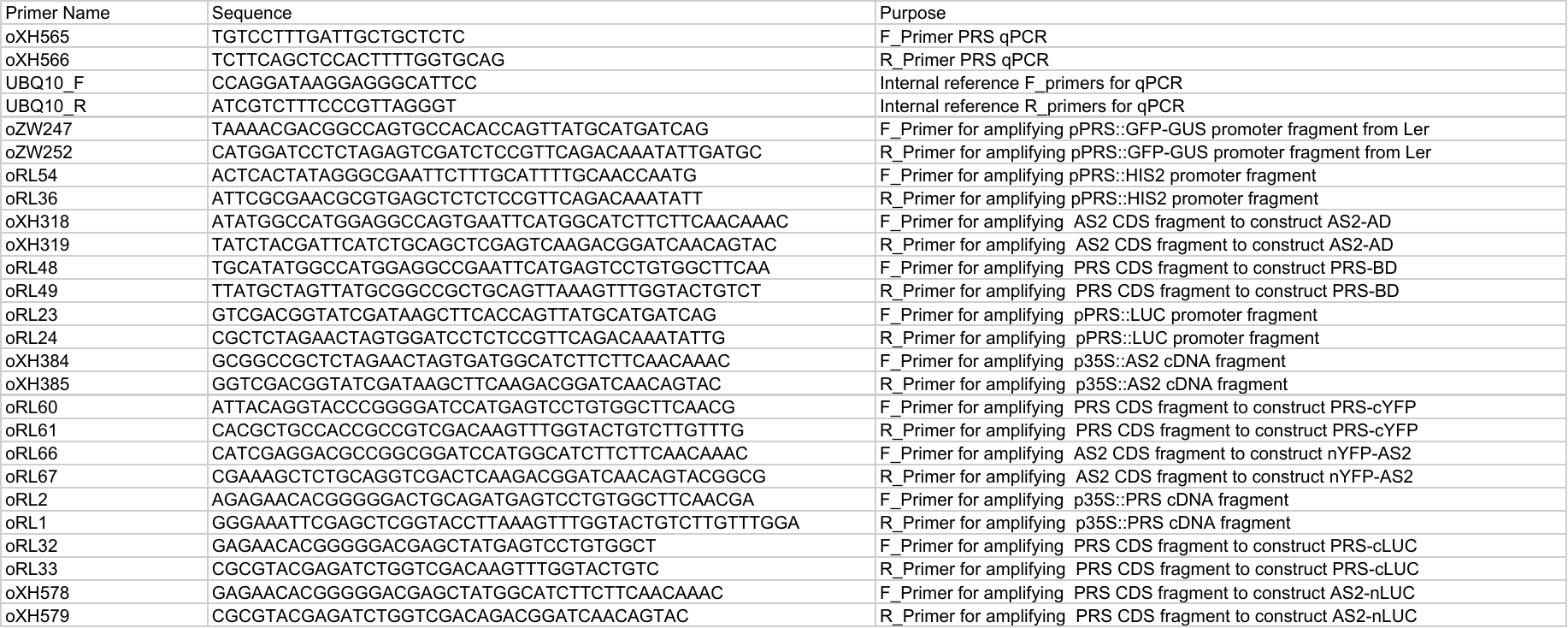
